# Normative Birth Weight Variation is Associated with White Matter Connectivity in Full-Term Neonates

**DOI:** 10.1101/2023.09.05.556390

**Authors:** Alexander J. Dufford, Wei Dai, Dustin Scheinost

## Abstract

**Importance:** Variation in birth weight, an indicator of fetal growth, has been prospectively associated with both alterations in brain development and developmental delays in cognitive and language domains. However, studies examining birth weight variation and white matter development in the brain typically test these associations in infants that are preterm or very low birth weight, leaving potential normative associations in full term infants unclear.

**Objective:** Here, we test prospective associations between birth weight variation in the ‘normative’ range (> 2.5 kg and <4.5 kg) and white matter connectivity in full-term neonates. Further, the main objective includes testing associations between normative birth weight variation and cognitive and language developmental scores at 18 months, and if white matter connectivity that is related to birth weight variation is further associated with cognitive language development. We hypothesized, greater normative birth weight would be associated with higher white matter connectivity controlling for gestational age, particularly in thalamic, inferior frontal, and middle temporal connectivity. Further, we predicted greater connectivity for these tracts would be associated with higher scores for cognitive (thalamic) and language development (inferior and middle temporal) at 18 months of age.

**Design:** The study was an observational longitudinal design of data from the Developing Human Connectome Project (dHCP). Birth weight was measured at the birth of the child, white matter connectivity was measured as neonates (mean=40.07 weeks, SD=1.14), and cognitive/language outcomes were measured at 18 months of age.

**Setting:** The dHCP data was collected at the Evelina Newborn Imaging Centre, Evalina London Children’s Hospital between 2015 and 2019.

**Participants:** A sub-sample of the full dHCP was tested. These participants were full-term neonates with birth weight variation in the ‘normative’ range (> 2.5 kg and < 4.5 kg). Participants also had to have usable diffusion-weighted imaging data as neonates and cognitive/language developmental assessments collected at 18 months (n=323).

**Exposure(s):** The study participants were all born at full-term and in the normative birth weight range.

**Main Outcome(s) and Measure(s):** The study had two main outcomes: white matter connectivity at the neonatal timepoint and cognitive/language developmental scores at 18 months. White matter connectivity was calculated from diffusion-weighted data for the whole-brain. Cognitive/language developmental scores were measured using the Bayley Scales of Infant and Toddler Development, Third Edition (Bayley-III) at 18 months.

**Results:** Using a Network-Based Statistic (NBS) approach, we found widespread associations between normative birth weight variation and white matter connectivity in full-term neonates, primarily in the positive direction for the right middle occipital gyrus and left supplementary motor area.

**Conclusions and Relevance:** While investigations have been focused on the extreme ends of the birth weight spectrum, we find evidence that there is a robust association between birth weight and white matter connectivity even within the normative birth weight range. As normative birth weight variation and regions of white matter associated with birth weight were further associated with language development scores at 18 months, our results suggest the birth weight to white matter pathway may be an underlying pathway between birth weight’s association with language development.

**Key Points:** *Question:* Are variations in normative birth weight associated with white matter connectivity and cognitive/language outcomes in infancy?

*Findings:* Greater normative birth weight is associated with greater white matter organization across a widespread network of connections in the neonatal brain. Greater white matter organization in this network for neonates has a positive prospective with expressive language development at 18 months of age.

*Meaning:* Even variations within the normative birth weight range have robust associations with early white matter development and can be prospectively linked to language development.

## Introduction

Birth weight is an indicator of fetal growth^1-4^. Low birth weight (LBW, below 2.5 kilograms) has a prospective association with various developmental challenges in cognition^5-7^ and communication/language^8-10^. The potential impact of birth weight on early brain development is hypothesized as a potential underlying pathway for these associations^5,7,8,11^. However, existing studies tend to focus on the tail of the birth weight spectrum—LBW and very LBW (VLBW; < 1.5 kg)—and include preterm (birth occurring before the 37^th^ week of pregnancy) infants. This focus limits the potential to examine which neurodevelopmental associations are common and unique between LBW/VLBW and within the normative range (> 2.5 kg and < 4.5 kg).

Several studies have identified associations between normative birth weight and developmental outcomes^12,13^. In a study of 467 mother-child dyads, lower birth weight in the normative range was associated with lower receptive language scores at 36 months^12^. In the cognitive domain, normative birth weight variation was positively associated with cognitive functioning at age 7 in a large, nationally representative sample of 3,484 children^14^. Therefore, even among normative birth weight, there is evidence of a prospective linear relationship with language and cognitive development.

Fewer studies have examined normative birth weight and brain development. In LBW and VLBW, converging evidence suggests lower birth weight is robustly associated with measures of white matter development. Early studies found evidence of myelination abnormalities in infants with VLBW^5,7,11^. These studies have been followed up with diffusion tensor imaging in adolescents and found lower fractional anisotropy (FA) in internal capsule, superior longitudinal, and inferior longitudinal fasciculi^11^. Lower FA in the internal capsule has been observed for infants that experienced VLBW^15^. However, associations between normative birth weight variation and white matter connectivity in neonates are unclear.

To address these gaps in knowledge, we leverage a large sample of full-term neonates from the Developing Human Connectome Project (dHCP) scanned close to birth with magnetic resonance imaging^16^. As studies have consistently found LBW to be associated with white matter development^5,8,15^, the present study examines associations between normative birth weight variation and the neonatal white matter connectome. We examined the prospective associations between birth weight variation in the ‘normative’ range and white matter connectivity, as measured by greater quantitative anisotropy (QA), in full-term neonates using data from the dHCP (n=323). We hypothesized that lower birth weight within the normative range would be associated with lower cognitive and language developmental scores at 18 months. We hypothesized that lower birth weight within the normative range would be associated with both lower cognitive and language developmental scores at 18 months of age. Further, we predicted greater connectivity in the inferior frontal and middle temporal white matter would be associated with higher scores for language development, and thalamic connectivity would be associated with higher cognitive development at 18 months. Lastly, as several perinatal exposures have been associated with birth weight and developmental outcomes^17-20^, we conducted an exploratory analysis to test if six types of perinatal exposures moderated associations between white matter connectivity and developmental outcomes. In addition, we conducted an exploratory mediation analysis to test if white matter connectivity mediated associations between birth weight and developmental outcomes.

## Materials and Methods

### Participants

Imaging and behavioral data were from the Developing Human Connectome Project (dHCP)^16^. The dHCP aims to understand human brain connectivity from 20 to 44 weeks post-conceptional age using neuroimaging, clinical, behavioral, and genetic data. Inclusion in the analytic sample required participants to have usable diffusion-weighted imaging data as neonates, have birth weight in the normative range, and developmental assessment at 18 months. This inclusion criteria resulted in an analytic sample of 323. The dHCP study was approved by the London Riverside Research Ethics Committee of the Health Research Agency. Written consent was obtained from each participating family before the study procedures. See the **Supplementary Information** for more information on the derivation of the analytic sample from the full sample. Demographic characteristics of the sample are presented in **Table 1**.

**Table 1.** Demographic characteristics of the sample.

| **Variable** | **Sample (N=323)** |
| --- | --- |
| Age at Birth (GA, months) | 40.1 (1.1) |
| Age at Scan (PMA, months) | 41.3 (1.6) |
| Sex (male) (1) | 171 (52.9%) |
| Birth Weight (kgs) | 3.4 (0.4) |
| Head Circumference at Scan (cm) | 34.1 (6) |
| Outlier Ratio from SHARD Scan | 0.2 (0.1) |
| Bayley-III Cognitive Composite Score | 100.7 (10.5) |
| Bayley-III Language Composite Score | 98.4 (15) |
| Receptive Language Subscale (Bayley) | 10.3 (3.1) |
| Expressive Language Subscale (Bayley) | 9.1 (2.6) |
| Age at Developmental Assessment (months) | 18.4 (1.6) |

### DWI Preprocessing

The full details of the DWI preprocessing are discussed in the **Supplementary Information**. Briefly, whole brain tracking was conducted using DSI Studio for the UNC Neonate Atlas.^21^ Tracking was conducted using quantitative anisotropy (QA). QA is like other measures of white matter organization, such as fractional anisotropy or FA, but has been shown to be less susceptible to noise, partial volume effects, and has increased spatial resolution for tractography^22^. Tracking using QA calculates the QA between node of the parcellation^22^. The preprocessing resulted in a 90x90 matrix for each neonate.

### Developmental Assessment at 18 Months

Cognitive and language development was assessed at a follow-up visit using the Bayley Scales of Infant and Toddler Development, Third Edition (Bayley-III)^23^. We focused on the cognitive and language domains as they have had the most consistent associations with normative and non-normative birth weight^6-10,12,24^. In addition to the focus on the cognitive composite and language composite scores, we conducted post-hoc sensitivity analyses to test if associations were specific to the expressive or receptive language subscales of the Bayley-III. The receptive subscale assesses the child’s ability to comprehend and respond to requests and discriminate between sounds in the environment^25^. The expressive subscale assesses the child’s ability to communicate wants, and name objects and actions.

### Network-based Statistic (NBS)

The NBS method^26^ is analogous to cluster-based correction used in fMRI activation-based studies but performed at the level of ‘groups’ of edges rather the cluster. We used two NBS models to examine associations between birth weight variation and white matter connectivity in which we examined both the positive and negative contrasts. The component-determining threshold was set at *z*=5.0 with 10,000 permutations and a *p*-value threshold of *p*<.05 corrected. Each model included participant’s gestational age, age at scan, sex, head circumference, and head motion as covariates.

### Associations with Developmental Assessments at 18 Months

For the NBS analysis, we extracted the connectivity values for the significant edges. We examined if the sum of QA from all the edges associated with birth weight were further associated with cognitive and language developmental scores. The extracted values were analyzed in R^27^ using a linear model to examine the association between the extracted values and language development scores (language compositive score of the Bayley-III) at 18 months. These linear models included gestational age, age at scan, sex, head circumference at scan, head motion. Multiple comparisons correction was performed using the Benjamini-Hochberg False Discovery Rate procedure at *q*<.05. As a post-hoc sensitivity analysis, we first tested if birth weight was associated with the receptive or expressive language subscales of the Bayley-III. We also examined if the sum of QA in all the edges associated with birth weight) was associated with receptive or expressive language scores at 18 months.

### Exploratory Moderation and Mediation Analyses

A full list of the six perinatal exposures tested and information about their measurement is presented in the **Supplementary Information**. As perinatal exposures are typically conceptualized as ‘moderators’, we examined if each variable moderated the association between white matter connectivity and language scores using multiple regressions adjusting for age at scan, gestational age at birth, sex, head circumference, and head motion.

A second exploratory analysis tested if white matter connectivity mediated associations between birth weight and developmental scores. We conducted post-hoc mediation models for analyses in which the requirements for mediation testing were met^28^ (associations between X, M, and Y variables). The mediation models were conducted using the “mediate” package in R^29^ with the *mediate* function. The indirect effect significance was tested with bootstrapping with 10,000 iterations. The mediation models included the same covariates as the linear and moderation models: age at scan, gestational age at birth, sex, head circumference, and head motion.

## Results

### Normative Birth Weight, Demographics, and Developmental Assessment Scores

The correlation between normative birth weight and cognitive and language developmental scores at 18 months was nonsignificant for the cognitive composite (*r*(321)= .06, *p*=0.22, 95% CI [-0.04,0.17]), and significant for language composite (*r*(321)= .12, *p*<.02, 95% CI [0.01, 0.23]). Therefore, we focused our subsequent analyses on the language scores at 18 months. Birth weight associated with receptive language was at a trend level (*r*(321)= .10, *p*<.05, 95% CI [-0.003, 0.21]) and significant for expressive language (*r*(321)= .12, *p*<.02, 95% CI [0.019, 0.23]). See **Table 2** for the correlation table of study variables.

**Table 2.**
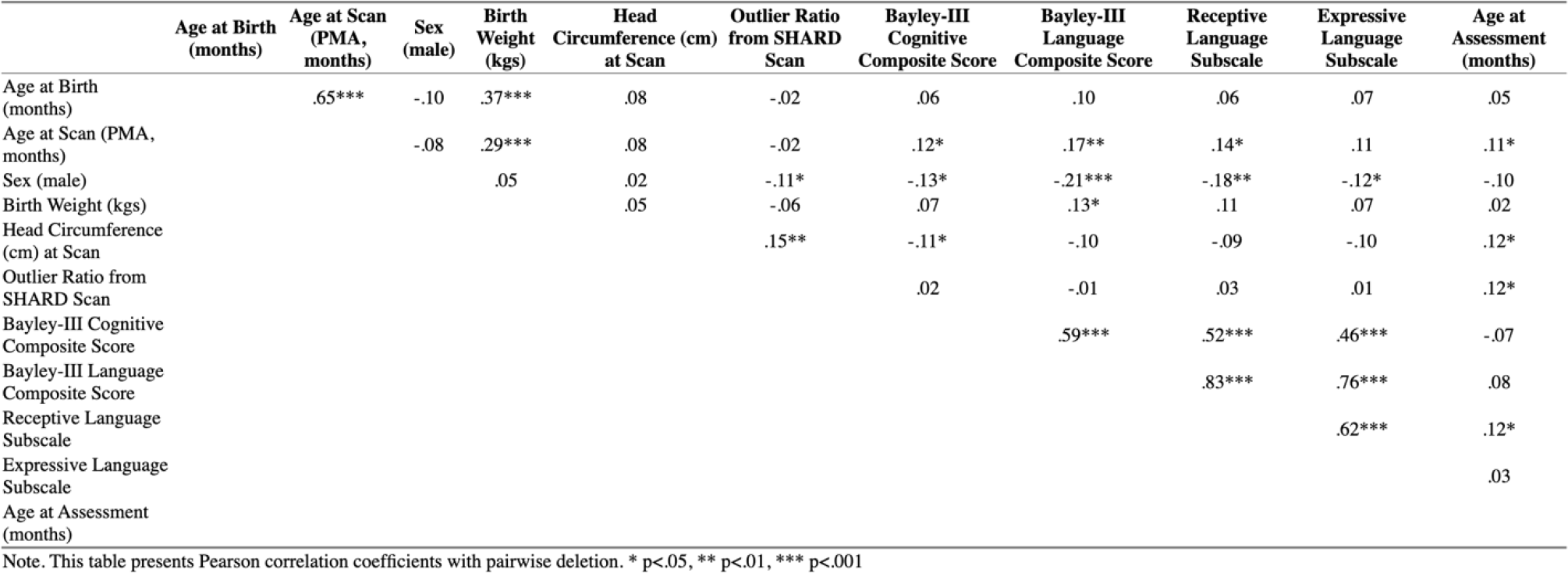
Correlation table for the study variables.

|  | Age at Birth<br>(months) | Age at Scan<br>(PMA,<br>months) | Sex<br>(male) | Birth<br>Weight<br>(kgs) | Head<br>Circumference (cm)<br>at Scan | Outlier Ratio<br>from SHARD<br>Scan | Bayley-III<br>Cognitive<br>Composite Score | Bayley-III<br>Language<br>Composite Score | Receptive<br>Language<br>Subscale | Expressive<br>Language<br>Subscale | Age at<br>Assessment<br>(months) |
| --- | --- | --- | --- | --- | --- | --- | --- | --- | --- | --- | --- |
| Age at Birth<br>(months) |  | .65*** | -.10 | .37*** | .08 | -.02 | .06 | .10 | .06 | .07 | .05 |
| Age at Scan (PMA,<br>months) |  |  | -.08 | .29*** | .08 | -.02 | .12* | .17** | .14* | .11 | .11* |
| Sex (male) |  |  |  | .05 | .02 | -.11* | -.13* | -.21*** | -.18** | -.12* | -.10 |
| Birth Weight (kgs) |  |  |  |  | .05 | -.06 | .07 | .13* | .11 | .07 | .02 |
| Head Circumference<br>(cm) at Scan |  |  |  |  |  | .15** | -.11* | -.10 | -.09 | -.10 | .12* |
| Outlier Ratio from<br>SHARD Scan |  |  |  |  |  |  | .02 | -.01 | .03 | .01 | .12* |
| Bayley-III Cognitive<br>Composite Score |  |  |  |  |  |  |  | .59*** | .52*** | .46*** | -.07 |
| Bayley-III Language<br>Composite Score |  |  |  |  |  |  |  |  | .83*** | .76*** | .08 |
| Receptive Language<br>Subscale |  |  |  |  |  |  |  |  |  | .62*** | .12* |
| Expressive Language<br>Subscale |  |  |  |  |  |  |  |  |  |  | .03 |
| Age at Assessment<br>(months) |  |  |  |  |  |  |  |  |  |  |  |

Note. This table presents Pearson correlation coefficients with pairwise deletion. \* p<.05, \*\* p<.01, \*\*\* p<.001

### Normative Birth Weight and White Connectivity Using NBS

The NBS model that test the positive association between normative birth weight and QA-based white matter connectomes was highly significant (*p*<.001, corrected) and widespread (see **Figure 1a**). It was comprised of 158 edges between 66 nodes. The most strongly associated edges were between the left olfactory cortex and left middle temporal gyrus (*t*=5.96), right superior occipital gyrus and right thalamus (*t*=5.86), right supramarginal gyrus and right middle temporal gyrus (*t*=5.74). The highest degree nodes were the right middle occipital gyrus (degree=12), left supplementary motor area (degree=12), right inferior temporal gyrus (degree=10). The negative contrast NBS model of birth weight variation and QA-based white matter connectomes was also highly significant (*p*<.001). The significant network was comprised of two nodes (left olfactory cortex and superior temporal pole) and 1 edge (*t*=5.11) (**Figure 2a**). The highest degree nodes were the right middle occipital gyrus (degree=12), left supplementary motor area (degree=12), right inferior temporal gyrus (degree=10). The negative contrast NBS model of birth weight variation and QA-based white matter connectomes was also highly significant (*p*<.001). The significant network was comprised of two nodes (left olfactory cortex and superior temporal pole) and one edge (*t*=5.11) (**Figure 2b**).

**Figure 1.**
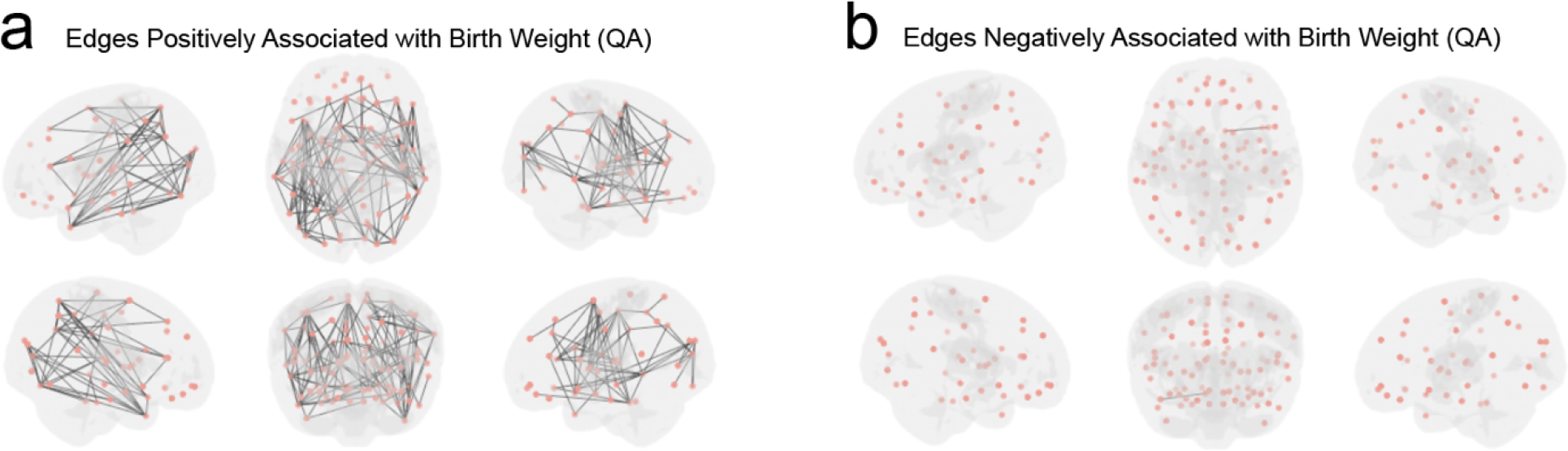
(a) Results of the Network-Based Statistic (NBS) analysis, indicating edges (white matter connectivity) positively associated with birth weight for quantitative anisotropy (QA). (b) Results for the negative contrast for associations with birth weight.

**Figure 2.**
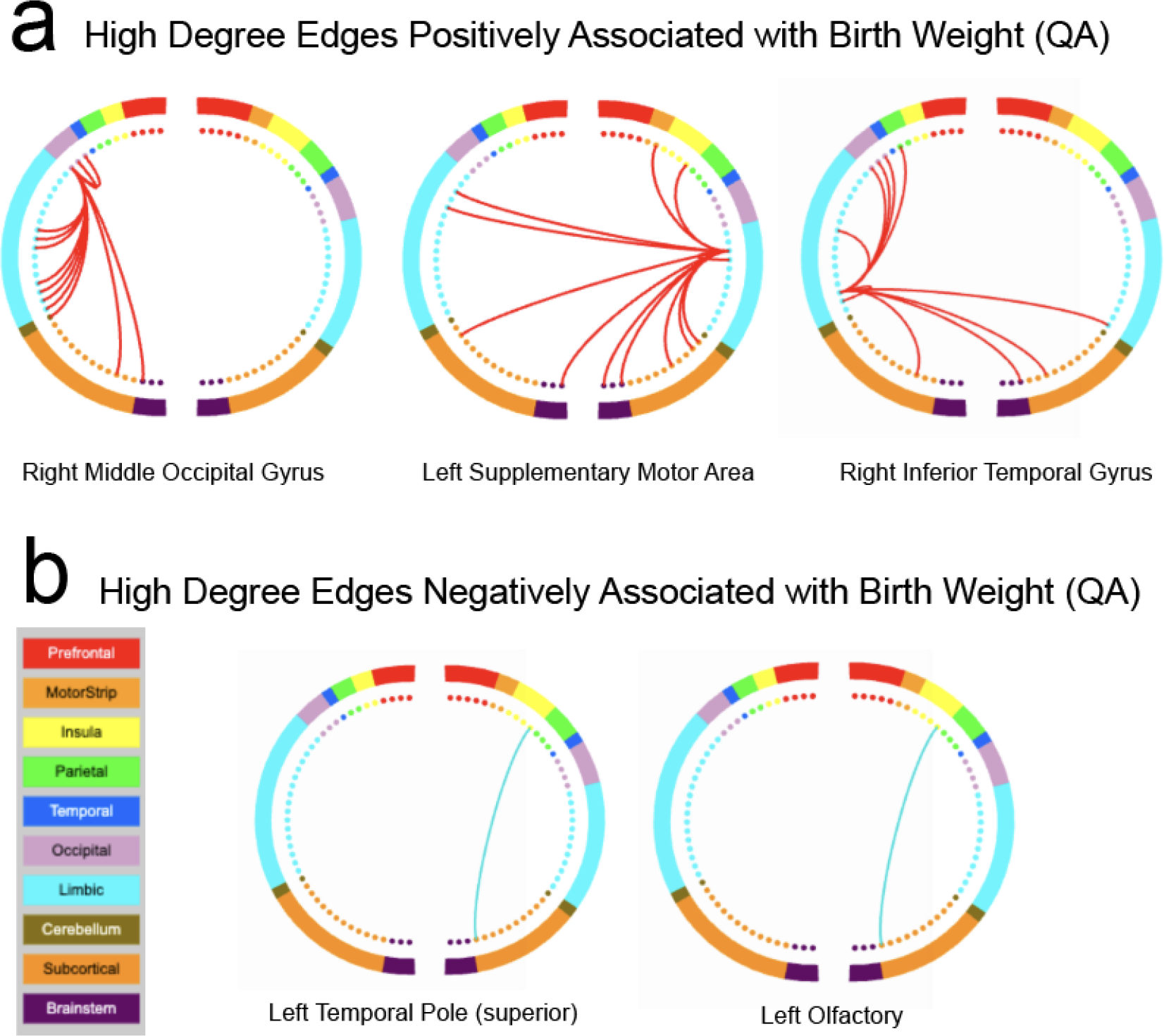
(a) Top three high degree nodes for the positive association between birth weight and quantitative anisotropy (QA) matrices. (b) High degree nodes for the negative association between birth weight and QA.

### Associations with Cognitive and Language Development Scores

Using the edges significantly correlated with birth weight from the NBS analysis, we summed the QA values across those edges to create a single summary score to associated with cognitive and language development scores. The regression equation testing the association between the sum of QA in the birth weight-related network and language scores was significant [(*F*(6, 316) = x, *p*<.001) with an *R*^*2*^ of .09 and Cohen’s *f*^2^ of .09)] (see **Supplementary Figure 1a**) and indicated an association at a trend level for QA and language scores (*b*=.002, *p*=.06, 95% CI [-0.0001, 0.006]). The association between the sum of QA in the birth weight-related network and cognitive development scores was nonsignificant (*p*=.89). As the birth weight-related network was widespread and included several edges, we conducted further analyses on the most significantly associated edges with birth weight. These results are presented in the **Supplementary Information**.

We conducted an additional post-hoc sensitivity analysis to examine if the associations between the extracted QA values (i.e., sum of QA in the birth weight-related network) were associated with receptive and/or expressive language scores (adjusting for age at scan, gestational age at birth, sex, head circumference, and head motion). The linear model for the sum of QA in the birth weight-related network association with expressive language scores was significant [(*F*(6,316) = 5.23, *p*<.05) with an *R*^*2*^ of .05 and Cohen’s *f*^2^ of .05)], indicating a significant positive association (*b*= 0.0006, *p*<.01, 95% CI [0.0001, 0.001]). The association between the sum of QA in the birth weight-related network and receptive language scores was nonsignificant (*p*<.19). Further the associations between the top 3 most significant edges and receptive/expressive language are presented in the **Supplementary Information**.

### Exploratory Analysis of Perinatal Exposures as Moderators

Of the six perinatal exposures tested, the only significant moderation effect was found for the depressive symptoms of the gestational parent (GP). Individuals that were exposure to a high level of perinatal depressive symptoms and had a low sum of QA in the birth weight-related network exhibited lower language scores at 18 months. For testing the moderation, the linear model was significant [(*F*(8,274) = 5.11, *p*<.001) with an *R*^*2*^ of .12 and Cohen’s *f*^2^ of .13)] and the sum of QA by depressive score interaction term was significant (*b*= 0.0007, *p*<.02, 95% CI [0.0001, 0.0014]). Perinatal exposures that were not moderators included history of diabetes for the GP, GP body mass index, and smoking during pregnancy (*ps*>.05), highlighting the robustness our results. The full details of this exploratory analysis are presented in the **Supplementary Information**.

### Exploratory Analysis of White Matter Connectivity as Mediator of Birth Weight Language Development Association

As an exploratory mediation analysis, we examined the indirect effect of birth weight on language development scores through the sum of QA in the birth weight-related network. The first mediation model tested the language composite scores as the outcome variable and the indirect effect was not significant (*p*=0.19, 95% CI [-0.34, 1.96]). However, the mediation model testing expressive language scores, the indirect effect was significant *(ab*=0.20, *p*=0.03, 95% CI [0.01, 0.44]) indicating sum of QA in the birth weight-related network as a significant mediator of the association between birth weight and expressive language scores

## Discussion

While associations between birth weight, white matter, and developmental outcomes have been reported^5,7^, few studies have focused on normative birth weight. Here, we found evidence of robust associations between normative birth weight and white matter connectivity in a sample of full-term infants. The findings supported our hypothesis that birth weight would be associated with middle temporal connectivity but less so for our hypothesis regarding inferior frontal connectivity. As birth weight has also been associated with developmental outcomes, we examined if regions of white matter connectivity associated with birth weight variation were further prospectively associated with language development at 18 months. The sum of QA in the birth weight-related network had a significant positive association with expressive language scores. The findings highlight the importance of considering variations in the normative birth weight range and further support the role of white matter development in birth weight and developmental outcome associations, specifically for expressive language.

Birth weight variation had a robust and widespread positive association with QA across the brain. This finding aligns with other studies of LBW and VLBW that found that higher birth weight is associated with higher anisotropy^8,11,30^. Regarding brain developmental processes, QA measures white matter organization, which can be influenced by axonal density and myelination^22^. Therefore, consistent with previous studies, lower birth weight may impact these processes, resulting in the lower white matter organization among these regions as neonates. While the QA results in the positive direction were widespread, consistent connectivity of the middle temporal gyrus was found for the most robust associations. Other studies of the low end of the birth weight spectrum found links with middle temporal connectivity, specifically for the superior longitudinal fasciculus (SLF) and inferior longitudinal fasciculus (ILF) in adoelescents^11^. We did not find support for our hypothesis that birth weight variation would be associated with thalamic connectivity. It may be that thalamic associations are unique to preterm birth^31-33^. The present study’s findings highlight the importance of considering the potential full spectrum of birth weight in terms of its associations with white matter connectivity.

The findings from the present study align with the theory of “developmental origins of health and disease” proposed by Barker^34^. This theory states that experience early in development (such as fetal growth) may have ‘programming’ effects on the developmental trajectories of an organism. Several studies have explained birth weight and developmental outcomes regarding the Barker hypothesis^35,36^. These myriad factors can potentially impact fetal growth and subsequent birth weight. They may have programming effects on white matter development (myelination, axonal maturation)^37-40^ and the development of language skills^9,24^.

Our findings highlight the importance of examining these factors as a continuous distribution across VLBW, LBW, normative weight, and LGA instead of only focusing on VLBW and LBW. Further, birth weight—as an indicator of fetal growth—may have advantages in accuracy as well-documented errors in gestational age estimation exist^41,42^.

As studies have hypothesized white matter development to be a potential underlying pathway of the birth weight to developmental outcome pathway, we examined if connectivity values associated with birth weight were further prospectively associated with language development scores at 18 months. Consistent with previous studies finding links between white matter connectivity associated with birth weight being further associated with language development^8^, participants with a greater sum of QA values in the birth weight-related network had higher expressive language scores. These findings align with previous studies that have found associations between the anisotropy, specifically the arcuate fasciculus/superior longitudinal fasciculus (AF/SLF), and language scores, including those in infancy^43-46^. When examining the top three most significantly birth weight-related edges, both connections involving the middle temporal gyrus (left olfactory cortex to left middle temporal gyrus and right supramarginal gyrus to right middle temporal gyrus connectivity) were associated with language scores (composite) and expressive language. Additionally, an exploratory mediation analysis found preliminary evidence of a mediating effect of the sum of QA in the birth weight-related network on the relation between birth weight and expressive language scores.

The present study has limitations that should be considered. First, the whole brain connectomic approach does not provide information about specific white matter tracts, the edges found in the present study can be used to generate hypotheses for analyses that use tractometry rather than connectomic approaches. Second, there are challenges and limitations associated with mediation analysis for neuroimaging data^47^. Therefore, the mediation findings are exploratory, should be interpreted with caution, and require replication in future studies.

## Conclusion

Using a large sample of term infants scanned as neonates, we found that greater normative birth weight was associated with greater language development scores at 18 months. This association was more robust in expressive versus receptive language. We found associations between normative birth weight and white matter connectivity, particularly in the positive direction for the white matter organization or QA between regions. The sum of QA in the birth weight-related network was not associated with cognitive development and a trend level for language developmental scores. Further, the sum of QA in the birth weight-related network was positively associated with expressive and not receptive language. The findings highlight the potential importance of variations within the normative birth weight range for developing white matter connectivity. Further, we demonstrate that QA related to birth weight had a prospective association with language development and may be an underlying pathway to explain birth weight-related variations in language ability.

## Data Statement

The raw data from this study can be accessed by the releases of the Developing Human Connectome Project (https://www.developingconnectome.org/data-release/third-data-release/) and the completion of a data use agreement. The preprocessed diffusion-weighted data for this study was downloaded from the DSI Studio webpage (https://brain.labsolver.org/hcp_d2.html).

## Data Availability

Links to download the data for the study have been provided in the Data Statement section.

## Acknowledgements

The authors would like to acknowledge and thank the Developing Human Connectome Project and the families that participated in the study.

## References

1 Lee, W. et al. Fetal growth parameters and birth weight: their relationship to neonatal body composition. Ultrasound in Obstetrics and Gynecology: The Official Journal of the International Society of Ultrasound in Obstetrics and Gynecology 33, 441–446 (2009).

2 Maulik, D. Fetal growth compromise: definitions, standards, and classification. Clinical obstetrics and gynecology 49, 214–218 (2006).

3 Wheater, E. et al. Birth weight is associated with brain tissue volumes seven decades later but not with MRI markers of brain ageing. NeuroImage: Clinical 31, 102776 (2021).

4 Rosenzweig, M. R. & Schultz, T. P. in Economic aspects of health 53-92 (University of Chicago Press, 1982).

5 Skranes, J. et al. White matter abnormalities and executive function in children with very low birth weight. Neuroreport 20, 263–266 (2009).

6 Shenkin, S. D., Starr, J. M. & Deary, I. J. Birth weight and cognitive ability in childhood: a systematic review. Psychological bulletin 130, 989 (2004).

7 Sato, J. et al. White matter alterations and cognitive outcomes in children born very low birth weight. NeuroImage: Clinical 32, 102843 (2021).

8 Reidy, N. et al. Impaired language abilities and white matter abnormalities in children born very preterm and/or very low birth weight. The Journal of Pediatrics 162, 719–724 (2013).

9 Byrne, J., Ellsworth, C., Bowering, E. & Vincer, M. Language development in low birth weight infants: the first two years of life. Journal of developmental and behavioral pediatrics: JDBP 14, 21–27 (1993).

10 Ribeiro, L. A. et al. Attention problems and language development in preterm low-birth-weight children: Cross-lagged relations from 18 to 36 months. BMC pediatrics 11, 1–11 (2011).

11 Skranes, J. et al. Clinical findings and white matter abnormalities seen on diffusion tensor imaging in adolescents with very low birth weight. Brain 130, 654–666 (2007).

12 Madigan, S., Wade, M., Plamondon, A., Browne, D. & Jenkins, J. M. Birth weight variability and language development: Risk, resilience, and responsive parenting. Journal of Pediatric Psychology 40, 869–877 (2015).

13 Wade, M., Browne, D., Madigan, S., Plamondon, A. & Jenkins, J. Normal birth weight variation and children’s neuropsychological functioning: Links between language, executive functioning, and theory of mind. Journal of the International Neuropsychological Society 20, 909–919 (2014).

14 Matte, T. D., Bresnahan, M., Begg, M. D. & Susser, E. Influence of variation in birth weight within normal range and within sibships on IQ at age 7 years: cohort study. Bmj 323, 310–314 (2001).

15 Dudink, J. et al. Fractional anisotropy in white matter tracts of very-low-birth-weight infants. Pediatric radiology 37, 1216–1223 (2007).

16 Edwards, A. D. et al. The developing human connectome project neonatal data release. Frontiers in neuroscience 16, 886772 (2022).

17 Chomitz, V. R., Cheung, L. W. & Lieberman, E. The role of lifestyle in preventing low birth weight. The future of children, 121-138 (1995).

18 Nigg, J. T. & Breslau, N. Prenatal smoking exposure, low birth weight, and disruptive behavior disorders. Journal of the American Academy of Child & Adolescent Psychiatry 46, 362–369 (2007).

19 Yang, Y. et al. The association of gestational diabetes mellitus with fetal birth weight. Journal of Diabetes and its Complications 32, 635–642 (2018).

20 Upadhyay, S., Biccha, R., Sherpa, M., Shrestha, R. & Panta, P. Association between maternal body mass index and the birth weight of neonates. Nepal Med Coll J 13, 42–45 (2011).

21 Shi, F. et al. Infant brain atlases from neonates to 1-and 2-year-olds. PloS one 6, e18746 (2011).

22 Yeh, F.-C., Verstynen, T. D., Wang, Y., Fernández-Miranda, J. C. & Tseng, W.-Y. I. Deterministic diffusion fiber tracking improved by quantitative anisotropy. PloS one 8, e80713 (2013).

23 Bayley, N. Bayley scales of infant and toddler development. (2006).

24 Barre, N., Morgan, A., Doyle, L. W. & Anderson, P. J. Language abilities in children who were very preterm and/or very low birth weight: a meta-analysis. The Journal of pediatrics 158, 766-774. e761 (2011).

25 Michalec, D. 215–215 (Boston, MA: Springer, 2011).

26 Zalesky, A., Fornito, A. & Bullmore, E. T. Network-based statistic: identifying differences in brain networks. Neuroimage 53, 1197–1207 (2010).

27 Ihaka, R. & Gentleman, R. R: a language for data analysis and graphics. Journal of computational and graphical statistics 5, 299–314 (1996).

28 MacKinnon, D. P., Fairchild, A. J. & Fritz, M. S. Mediation analysis. Annu. Rev. Psychol. 58, 593–614 (2007).

29 Revelle, W. & Revelle, M. W. Package ‘psych’. The comprehensive R archive network 337 (2015).

30 Eikenes, L., Løhaugen, G. C., Brubakk, A.-M., Skranes, J. & Håberg, A. K. Young adults born preterm with very low birth weight demonstrate widespread white matter alterations on brain DTI. Neuroimage 54, 1774–1785 (2011).

31 Ball, G. et al. The effect of preterm birth on thalamic and cortical development. Cerebral cortex 22, 1016–1024 (2012).

32 Ball, G. et al. The influence of preterm birth on the developing thalamocortical connectome. Cortex 49, 1711–1721 (2013).

33 Menegaux, A. et al. Impaired visual short-term memory capacity is distinctively associated with structural connectivity of the posterior thalamic radiation and the splenium of the corpus callosum in preterm-born adults. Neuroimage 150, 68–76 (2017).

34 Godfrey, K. M. & Barker, D. J. Fetal programming and adult health. Public health nutrition 4, 611–624 (2001).

35 Reyes, L. & Manalich, R. Long-term consequences of low birth weight. Kidney International 68, S107–S111 (2005).

36 Daniels, S. R. The Barker hypothesis revisited. The Journal of Pediatrics 173, 1–3 (2016).

37 Miller, S. L., Huppi, P. S. & Mallard, C. The consequences of fetal growth restriction on brain structure and neurodevelopmental outcome. The Journal of physiology 594, 807–823 (2016).

38 Dudink, I. et al. Altered trajectory of neurodevelopment associated with fetal growth restriction. Experimental neurology 347, 113885 (2022).

39 Noma, K., Tamai, H., Shimada, S. & Funato, M. Myelination in very low birth weight infants--evaluation by MRI. No to Hattatsu= Brain and Development 23, 336–341 (1991).

40 Skranes, J. S., Nilsen, G., Smevik, O., Vik, T. & Brubakk, A.-M. Cerebral MRI of very low birth weight children at 6 years of age compared with the findings at 1 year. Pediatric radiology 28, 471–475 (1998).

41 Skupski, D. W. et al. Estimating gestational age from ultrasound fetal biometrics. Obstetrics and gynecology 130, 433 (2017).

42 Robson, S. C., Gallivan, S., Walkinshaw, S. A., Vaughan, J. & Rodeck, C. H. Ultrasonic estimation of fetal weight: use of targeted formulas in small for gestational age fetuses. Obstetrics & Gynecology 82, 359–364 (1993).

43 Zuk, J. et al. White matter in infancy is prospectively associated with language outcomes in kindergarten. Developmental Cognitive Neuroscience 50, 100973 (2021).

44 Dick, A. S. & Tremblay, P. Beyond the arcuate fasciculus: consensus and controversy in the connectional anatomy of language. Brain 135, 3529–3550 (2012).

45 Huber, E., Corrigan, N. M., Yarnykh, V. L., Ramårez, N. F. & Kuhl, P. K. Language experience during infancy predicts white matter myelination at age 2 years. Journal of Neuroscience 43, 1590–1599 (2023).

46 Salvan, P. et al. Language ability in preterm children is associated with arcuate fasciculi microstructure at term. Human Brain Mapping 38, 3836–3847 (2017).

47 Kriegeskorte, N., Simmons, W. K., Bellgowan, P. S. & Baker, C. I. Circular analysis in systems neuroscience: the dangers of double dipping. Nature neuroscience 12, 535–540 (2009).

