## Supplementary material for "Normative Birth Weight Variation is Associated with White Matter Connectivity in Full-Term Neonates": [Supplementary Information]

**Supplementary Methods**

**Participants.** The objectives and descriptions of the dataset and data releases of dHCP have been described elsewhere (<http://www.developingconnectome.org>). The dHCP study was approved by the London­ Riverside Research Ethics Committee of the Health Research Agency. Written consent was obtained from each participating family before the study procedures. Neonates were scanned at the Evelina Newborn Imaging Centre, Evalina London Children’s Hospital between 2015 and 2019. The present study used data from the dHCP’s Third Data Release of images from 783 neonatal participants.

Of the 724 participants’ DWI data that passed quality control, we removed repeated scans (as only a subset of the dataset had two scan timepoints) resulting in 624 participants with usable DWI data. To avoid confounds related to multiple births, we removed all DWI data from twin ‘B’ and kept all DWI data for twin ‘A’ resulting in 599 participants. Of these 599 participants, 460 had usable DWI data and language development data at 18 months. For these 460 participants, 112 were considered preterm defined as a birth age < 37 weeks. Due to the focus on examining birth weight variation in isolation of preterm birth, these 112 participants were removed. Of the remaining 348 full term neonates, 3 were removed due to a failed DWI reconstruction and 2 additional neonates were removed due failure of the tracking algorithm to successfully conduct the whole brain tractography. The final analytic sample consisted of 343 full-term neonates, that had whole brain tractography data that passed quality control procedures and had language developmental assessments at 18 months of age. We removed 15 additional participants because their birth weights were less than 2.5 grams and therefore would be considered ‘low birth weight’. Additionally, we removed 5 participants whose birth weight would be considered LGA (greater than 4.5 kg). These exclusions resulted in a final analytic sample of 323 participants.

**DWI Acquisition and Quality Control.** DWI data were acquired as a part of a larger neuroimaging protocol for the dHCP. Data were acquired using a 3-Telsa Philips Achieva system (Philips Medical Systems) with a dedicated 32-channel neonatal head coil. The preparation and scanning details have been previously described^13^. This procedure including no use of sedation, neonates were scanned during their natural sleep, with ear putty and earmuffs for additional attenuation of scanner noise. Each neonate was fed, swaddled, and placed in a vacuum jacked to promote continued sleep and a neonatal nurse and/or pediatrician monitored the heart rate, oxygen saturation, and temperature throughout the scan.

Preprocessed data was downloaded from <https://brain.labsolver.org/hcp_d2.html>. A multi-shell diffusion scheme was used with the following b-values were 400, 1000, and 2600 s/mm². The number of diffusion sampling directions were 64, 88, and 128, respectively. The in-plane resolution was 1.5 mm. The slice thickness was 1.5 mm. The images were denoised and corrected for Gibbs ringing, motion, eddy current, and susceptibility artifact using the diffusion SHARD pipeline. A quality check was conducted using neighboring DWI correction (NDC)^20^. Thirty-four out of 758 scans (including repeated scans) were excluded due to their low NDC values identified by a median value-based outlier detector. The accuracy of b-table orientation was examined by comparing fiber orientations with those of a population-averaged template^21^.

The restricted diffusion was quantified using restricted diffusion imaging^20^. The diffusion data were reconstructed using generalized q-sampling imaging^21^ with a diffusion sampling length ratio of 1.25 and tensor metrics were calculated. The analysis was conducted using the resource allocation (TG-CIS200026) at Extreme Science and Engineering Discovery Environment (XSEDE) resources^22^. For the analytic sample, white matter connectomes were calculated using Whole Brain Fiber Tracking in DSI Studio^20^ using the recommended settings (fiber count = 100000); however, the minimum length parameters were lowered to 20 due to tracking being for the neonatal brain. Tracking was conducted using the UNC Neonatal AAL atlas^21^ to calculate the ‘connectivity’ between each of its 90 nodes. The analysis focused on calculating the ‘quantitative anisotropy’ or QA between nodes^22^. QA is like other measures of white matter organization, such as fractional anisotropy or FA, but has been shown to be less susceptible to noise and has increased spatial resolution for tractography^22^. QA, like FA, is bounded between 0 and 1 where greater values indicate ‘greater’ white matter organization. QA’s calculation has been previously described. As a measure of head motion during the scan we used the slice ‘outlier ratio’. This metric measures the number of slicer outliers based upon signal dropout or artefact due to bulk motion and is calculated during the slice-to-volume reconstruction of the SHARD data^23^.

**Network-Based Statistic Analysis.** For the Network-Based Statistic, a component-determining threshold is set and suprathreshold links are constructed based upon this threshold and any connected structures or ‘components’ are identified. Permutation testing is used to create a null distribution for the expected component size due to chance by shuffling the correspondence between the connectome and behavioral data. These are used to calculate *p*-values for each identified ‘component’ by comparing its size to a null distribution of maximal component size. Therefore, the family-wise error corrected *p*-value for a component size is calculated as the proportion of permutations with components of equal or larger size^26^.

**Main Effects of the Covariates.** To demonstrate the robustness of the birth weight association with white matter connectivity, we used NBS models to test for a main effect of the covariates in the birth weight analysis. This included examining the main effect for age at MRI scan (adjusting for gestational age at birth, sex, head circumference, head motion, and birth weight), gestational age at birth, sex, head circumference, and head motion. The NBS models were identical to the main birth weight NBS model in terms of settings with the component-determining threshold was set at z=5.0 with 10,000 permutations and we examined both the positive and negative contrasts. For these analyses, we also identified edges that were both significant for the main effect of the covariate and significantly associated with birth weight.

**Developmental Assessment at 18 Months.** Of the full sample of neonates scanned for the dHCP, 619 (79%) participated in a follow-up visit that was planned for 18 months corrected age. Due to disruptions from the COVID-19 pandemic, some participants were not able to attend, or assessments were slightly delayed (median assessment of 18 months + 12 days).

**Perinatal Exposures.** The six perinatal exposures tested for a moderation of the association between white matter connectivity and developmental scores were: history of diabetes for the gestational parent (GP), body mass index at the time of the scan for the GP, history of hypertension for the GP, smoking during the pregnancy, alcohol use during the pregnancy, pre-pregnancy weight, and depressive symptoms, as measured by the EPDS, at the time of the scan. While self-reported drug exposure was collected for the study, in our sub-sample only one GP reported drug use during pregnancy, and we were unable to test the moderation.

**Supplementary Results**

**Main Effects of the Covariates.** The NBS models for the main effects of sex, head circumference, and head motion were all nonsignificant (*ps*>.05). The positive association between gestational age at birth was significant (*p*<.001) with one edge between two nodes (*t*=5.2). The significant positive edge for gestational age at birth was between the left anterior cingulate and left posterior cingulate gyrus. No edges in the negative direction were significant for the association with gestational age at birth.

The main effect of age at scan in the positive direction for white matter connectivity was highly significant (*p*<.001). It was comprised of 751 edges between 88 nodes. The strongest associations were found between the left orbitofrontal cortex (middle) and right orbitofrontal cortex (superior) (*t*=8.44), right cuneus and right inferior temporal gyrus (*t*=8.48) and left orbitofrontal cortex (middle) and right orbitofrontal cortex (middle) (*t*=8.66). The highest degree nodes were found in the right orbitofrontal cortex (degree=40), left supplementary motor area (degree=37), and left parahippocampal gyrus (degree=35). The negative association between age at MRI scan and white matter connectivity was also highly significant (*p*<.001) but less widespread. It was comprised of 28 edges between 24 nodes. The edges with the strongest associations were between the left posterior cingulate gyrus and left pallidum (*t*=6.72), right posterior cingulate gyrus and left putamen (*t*=6.76) and left posterior cingulate gyrus and left putamen (*t*=6.84). High degree nodes for the negative association between age at scan and QA edges were found in the right gyrus rectus (degree=10), left dorsal superior frontal gyrus (degree=8), and right putamen (degree=5).

**Overlapping Edges between Birth Weight and Age at Scan.** In terms of overlapping edges, 147 edges were both positively associated with both age at birth and birth weight (see **Supplementary Figure 2**) and high degree nodes were in the right middle occipital gyrus (degree=11), left supplementary motor area (degree=10), and right inferior temporal gyrus (degree=10). There were no edges that were both negatively associated with age at scan and birth weight.

**Associations with Cognitive and Language Development Scores.** We extracted from the top three edges that were significantly associated with birth weight: connectivity between the left olfactory cortex and left middle temporal gyrus, right superior occipital gyrus and right middle temporal gyrus, and right supramarginal gyrus and right middle temporal gyrus. The model for left olfactory cortex to left middle temporal gyrus connectivity was significant (*F*(6,316) = 5.62, *p*<.001) with an *R^2^* of .09 and Cohen’s *f*^2^ of .09. Greater QA values between these regions was associated with greater language scores at 18 months (*b*=33.83, *p*<.01, 95% CI [5.58, 62.08]) (see **Supplementary Figure 1b**). QA values between the right superior occipital gyrus and right middle temporal gyrus were not associated with language scores at 18 months (*p*=0.14, 95% CI [-7.02, 45.9]). The regression was significant for the right supramarginal gyrus and right middle temporal gyrus model (*F*(6,316) = 5.56, *p*<.02) with an *R^2^* of .09 and Cohen’s *f*^2^ of .09. QA values between these regions were associated with greater language scores (*b*=33.83, *p*<.01, 95% CI [9.61, 127.46]) (see **Supplementary Figure 1c**). These associations survived FDR correction (both at *q*<.03). The extracted values for the edge that was negatively associated with birth weight for QA (left olfactory cortex to superior temporal pole connectivity), was not associated with language scores (*p*=0.53, 95% CI [-91.65, 47.79]).

**Perinatal Exposures.** For the exploratory analysis of perinatal exposures as moderators of the association between white matter connectivity and language developmental scores, only depressive symptoms were significant. For the EPDS higher scores indicate higher depressive symptoms with a cut-off score of 11 shown to maximize both sensitivity and specificity^29^. Forty-one participants of the 323 did not have EPDS data and were removed from this analysis. We did not find significant moderation for history of diabetes for the gestational parent (GP) (*p*=.78), body mass index at the time of the scan for the GP (*p*=.35), smoking during the pregnancy (*p*=.86), alcohol use during the pregnancy (*p*=.36), and pre-pregnancy weight (*p*=.81). **Supplementary Figure 3** shows the QA by depressive symptoms interaction for language development scores.

**Supplementary Figures**

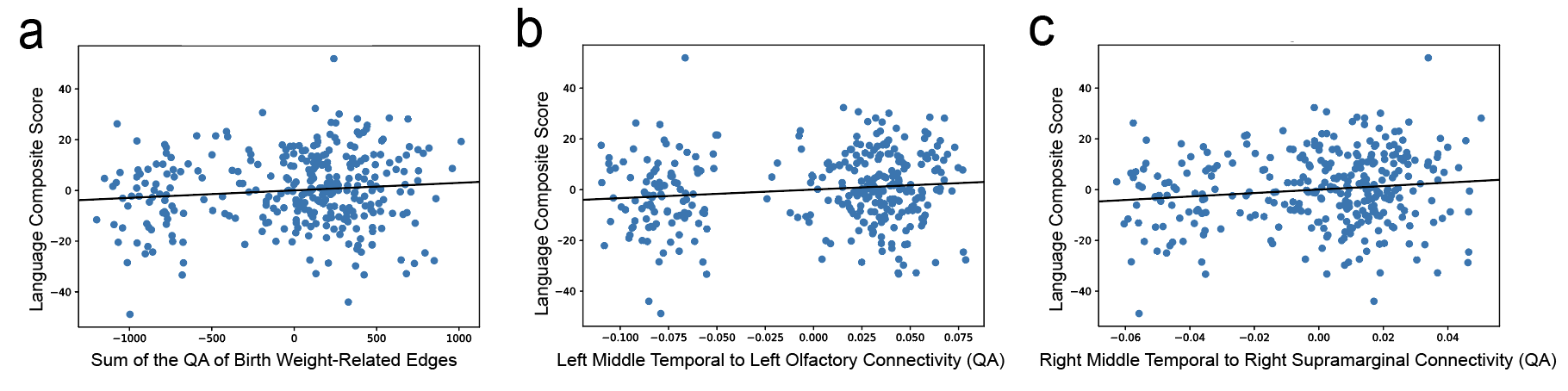

**Supplementary Figure 1.** (a) Partial regression plot of the association between QA of the left middle temporal to left olfactory edge and language scores from the developmental assessment at 18 months of age. (b) Partial regression plot of the association between QA of the left middle temporal gyrus to the left olfactory cortex and language scores. (c) Partial regression plot of the association between QA of the right middle temporal gyrus to right supramarginal gyrus connectivity and language scores. *All the plots were adjusted for age at MRI scan, gestational age at birth, sex, head circumference (at scan) and head motion.

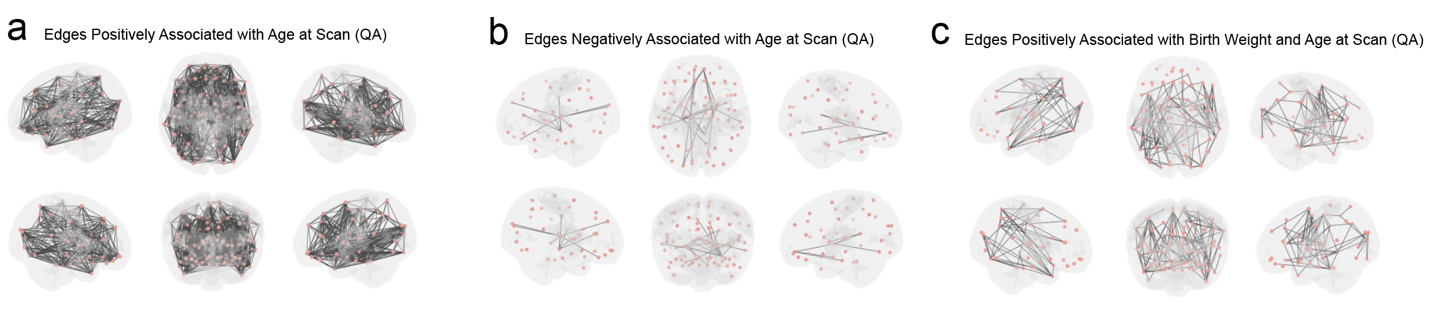

**Supplementary Figure 2.** (a) Results of the Network-based Statistic (NBS) model testing for the main effect of age at MRI scan on white matter connectivity, in the positive direction, as measured by quantitative anisotropy (QA). (b) Results of the Network-based Statistic (NBS) model testing for the main effect of age at MRI scan on white matter connectivity, in the negative direction, as measured by QA. (c) Edges that were both positively associated with birth weight and age at MRI scan.

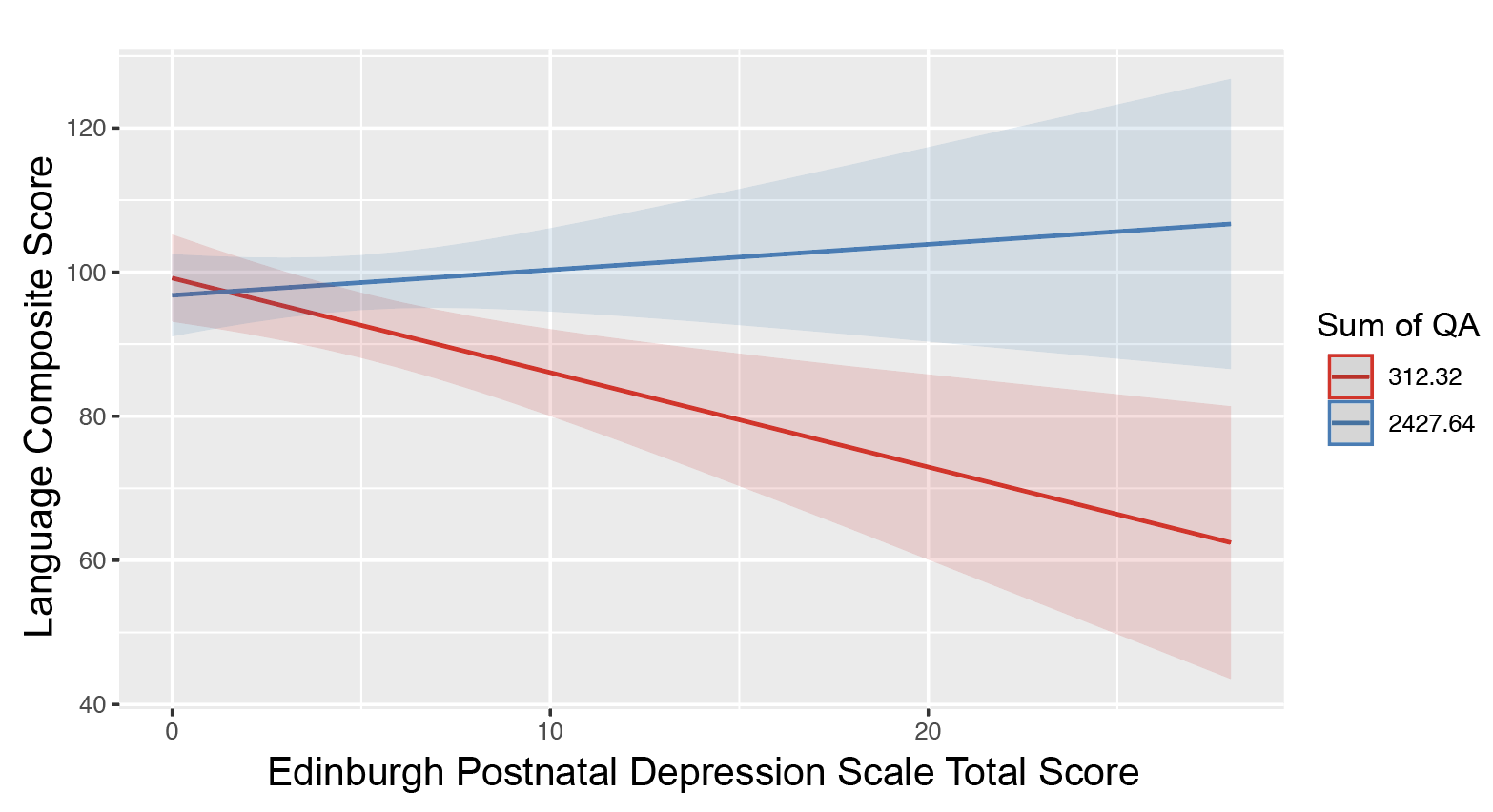

**Supplementary Figure 3.** Decomposition of the interaction of depressive scores (EPDS) and sum of the QA of birth weight-related edges for language development scores at 18 months. Participants with exposure to high depressive symptoms for their gestational parent and lower QA across the birth weight-related networks had lower scores on the language developmental assessment.
